# SARS-CoV-2 infects lung epithelial cells and induces senescence and an inflammatory response in patients with severe COVID-19

**DOI:** 10.1101/2021.01.02.424917

**Authors:** Konstantinos Evangelou, Dimitris Veroutis, Periklis G. Foukas, Koralia Paschalaki, Nefeli Lagopati, Marios Dimitriou, Angelos Papaspyropoulos, Orsalia Hazapis, Aikaterini Polyzou, Sophia Havaki, Athanassios Kotsinas, Christos Kittas, Athanasios G. Tzioufas, Laurence de Leval, Demetris Vassilakos, Sotirios Tsiodras, Ioannis Karakasiliotis, Peter J Barnes, Vassilis G. Gorgoulis

## Abstract

**Rationale:** SARS-CoV-2 infection of the respiratory system can progress to a life threatening multi-systemic disease, mediated via an excess of cytokines (“cytokine storm”), but the molecular mechanisms are poorly understood.

**Objectives:** To investigate whether SARS-CoV-2 may induce cellular senescence in lung epithelial cells, leading to secretion of inflammatory cytokines, known as the senescence-associated secretory phenotype (SASP).

**Methods:** Autopsy lung tissue samples from eleven COVID-19 patients and sixty age-matched non-infected controls were analysed by immunohistochemistry for SARS-CoV-2 and markers of cellular senescence (SenTraGor, p16^INK4A^) and key SASP cytokines (interleukin-1β, interleukin-6). We also investigated whether SARS-CoV-2 infection of an epithelial cell line induces senescence and cytokine secretion.

**Measurements and Main Results:** SARS-CoV-2 was detected by immunocytochemistry and electron microscopy predominantly in alveolar type-2 (AT2) cells, which also expressed the angiotensin-converting-enzyme 2 (ACE2), a critical entry receptor for this virus. In COVID-19 samples, AT2 cells displayed increased markers of senescence [p16^INK4A^, SenTraGor staining positivity in 12±1.2% of cells compared to 1.7±0.13% in non-infected controls (p<0.001)], with markedly increased expression of interleukin-1β and interleukin-6 (p<0.001). Infection of epithelial cells (Vero E6) with SARS-CoV-2 *in-vitro* induced senescence and DNA damage (increased SenTraGor and γ-H2AX), and reduced proliferation (Ki67) compared to uninfected control cells (p<0.01).

**Conclusions:** We demonstrate that in severe COVID-19 patients, AT2 cells are infected with SARS-CoV-2 and show senescence and expression of proinflammatory cytokines. We also show that SARS-CoV-2 infection of epithelial cells may induce senescence and inflammation, indicating that cellular senescence may be an important molecular mechanism of severe COVID-19.

## Introduction

The severe acute respiratory syndrome coronavirus 2 (SARS-CoV-2) causes the Coronavirus disease 2019 (COVID-19) that primarily affects the respiratory system. The clinical course of the patients ranges from asymptomatic to a life-threatening respiratory failure accompanied by a multi-systemic inflammatory disease (1,2). Systemic disease may occur through a viral-mediated “cytokine storm” that consists of a variety of cytokines and chemokines (CXCL-10, CCL-2, IL-6, IL-8, IL-12, IL1β, IFN-γ, TNF-α) (3,4). The link between viral infection of cells and development of severe lung disease and systemic manifestations is still poorly understood. Viral infection results in the activation of complex innate and adaptive immune responses that are orchestrated sequentially, involving several cell types and inflammatory mediators (5,6). At the cellular level, intrinsic defence mechanisms are activated and outcomes range from complete recovery to cell death (7–11). An “intermediate” and essential cellular state that is overlooked, due to lack of efficient methodological tools, is cellular senescence (12,13).

Cellular senescence is a stress response mechanism that preserves organismal homeostasis. Senescent cells are characterized by prolonged and generally irreversible cell-cycle arrest and resistance to apoptosis (12,14). Additionally, they also exhibit secretory features collectively described, as the senescence-associated secretory phenotype (SASP) (12). SASP includes a variety of cytokines, chemokines, growth factors, proteases and other molecules, depending on the senescence type (12,15). They are released in the extracellular space as soluble factors, transmembrane proteins following ectodomain shedding, or as molecules engulfed within small exosome-like vesicles (16–18). Under physiological conditions, senescence is transiently activated and SASP mediates the recruitment of immune cells for senescent cell clearance. In addition, other SASP factors promote tissue regeneration and repair, overall ensuring cellular/tissue homeostasis. On the contrary, persistence of senescent cells exerts harmful properties promoting tissue dysfunction and the maintenance of a “latent” chronic inflammatory milieu, via paracrine and systemic SASP (12,15).

There is little published evidence linking viral infection to cellular senescence (19–22). Given the significance of the “cytokine storm” in the progression of COVID-19 and the SASP secretion by senescent cells, we investigated whether cellular senescence occurs in COVID-19. We provide the first evidence supporting not only the evidence for senescence in COVID-19 infected lung cells, but also potential long-term adverse implications of this disease process.

## Materials and Methods

### Lung tissue

Formalin Fixed and Paraffin embedded autopsy lung tissue samples from eleven patients that died from COVID-19 (confirmed by RT-qPCR) and lung tissues resected prior to the COVID-19 outbreak, comprising a cohort of sixty previously published and new cases (negative controls) were analyzed (**Suppl. Table 1**) (23). Clinical sample collection and their experimental use were approved by the Commission Cantonale D’éthique de la Recherche, University of Lausanne, Switzerland (Ref 2020-01257), the Bio-Ethics Committee of University of Athens Medical School, Greece.

### Anti-SARS-COV-2 (G2) antibody generation

Mice immunization and antibodies collection, selection and specificity determination are described in detail in **Suppl Information.** Transcriptome analysis of hybridomas and amino acid determination of selected clones are also provided in **Suppl Information.** Four clones, namely 479-S1, 480-S2, 481-S3 and 482-S4 are under patent application **(Gorgoulis V.G., Vassilakos D. and Kastrinakis N. (2020) GR-patent application no: 22-0003846810).**

### Cells and SARS-CoV-2 culture

SARS-CoV-2 [isolate 30-287 (B.1.222 strain)] was obtained through culture in Vero E6 cells (ATCC^®^ CRL-1586), from an infected patient in Greece. The virus was recovered from a nasopharyngeal swab, rinsed in 1 ml saline and filtered twice through a 0.22 nm filter. Virus stock was prepared by infecting fully confluent Vero E6 cells in DMEM, 10% fetal bovine serum (FBS), with antibiotics, at 37°C, 5% CO_2_. Virus stock was collected four days after inoculation, sequenced by NGS (**Suppl Information**) and the supernatant was frozen (−80°C) until use. Infections were carried out in 24-well plates, using SARS-CoV-2 at a 0.01 MOI. Cells were either fixed with 4% paraformaldehyde or lysed with NucleoZOL (MACHEREY-NAGEL) 17 days post infection. Manipulations were carried out in a Biosafety level 3 facility.

**RNA extraction and Reverse-Transcription real-time PCR (RT-qPCR)** detection were performed as previously described (**Suppl Information**) (24).

### Next Generation Sequencing (NGS)

NGS was performed as previously described (25). Briefly, the Ion AmpliSeq Library Kit Plus was used to generate libraries following the manufacturer’s instruction, employing the Ion AmpliSeq SARS-CoV-2 RNA custom primers panel (ID: 05280253, Thermo Fisher Scientific). Briefly, library preparation steps involved reverse transcription of RNA using the SuperScript VILO cDNA synthesis kit (Thermo Fisher Scientific), 17-19 cycles of PCR amplification, adapter ligation, library purification using the Agencourt_AMPure XP (Beckman Coulter), and library quantification using Qubit Fluorometer high-sensitivity kit. Ion 530 Chips were prepared using Ion Chef and NGS reactions were run on an Ion GeneStudio S5, ion torrent sequencer (Thermo Fisher Scientific). Samples were run in triplicates.

### Immunocytochemistry and Immunohistochemistry

ICC and IHC were performed according to published protocols (26). The following primary antibodies were applied overnight at 4°C: i) anti-SARS-CoV-2 (G2) monoclonal antibody (at a dilution 1:300), ii) anti-ACE-2 (Abcam), iii) anti-TTF-1 (Dako), iv) anti-CD68 (Dako) and v) anti-p16^INK4A^ (Santa Cruz), vi) anti-IL-1β (Abcam) and vii) anti-IL-6 (R&D systems), viii) anti- phospho-histone (Ser 139) 2AX (γH2AX) (Cell Signaling) and ix) anti-Ki67 (Abcam)(**Suppl Information**).

**SenTraGor™ staining and double staining** experiments were performed and evaluated as previously described (26).

### Electron Microscopy

Representative area from hematoxylin and eosin stained paraffin sections of the lung autopsy of COVID-19 patients and corresponding non COVID-19 controls were chosen under the light microscope and marked. Paraffin-embedded tissue was deparaffinized, rehydrated and fixed in 2.5% glutaraldehyde in PBS for 24h and post-fixed in 1% aqueous osmium tetroxide for 1h at 4°C. The tissue fragment was embedded in fresh epoxy resin mixture, stained with ethanolic uranyl acetate and lead citrate and observed with a FEI Morgagni 268 transmission electron microscope equipped with Olympus Morada digital camera.

### Statistical analysis

The Wilcoxon paired non-parametric test was used to compare GL13 labelling indices and levels of IL-6, IL-8 and IL-1β between two groups (non-COVID-19 and COVID-19 infected).

## Results

### Detection of SARS-CoV-2 in lung cells

In order to detect SARS-CoV2 in lung tissue we developed monoclonal antibodies which react against the spike protein of SARS-CoV-2 and identified a high affinity antibody (G2) (**Suppl Figure 1, Suppl Figure 2, Suppl. Table 1A and Suppl. Table 1B)** (23). SARS-CoV-2 was detected predominantly in alveolar type 2 (AT2) cells, which are identified by TTF-1 positivity, and in sparse inflammatory cells (alveolar and tissue macrophages) in all COVID-19 patients (**Figure 1A, C**), ranging from <5 cells/4mm^2^ tissue to >50 cells/4mm^2^ tissue (**Suppl. Table 1A**). SARS-CoV-2 infected AT2 cells were occasionally large and appeared isolated (denuded or syncytial) or clustered (hyperplasia), exhibiting a variety of topological distribution (**Figure 1A**). These cells co-expressed the angiotensin-converting enzyme 2 (ACE2) receptor (**Figure 1B**), supporting SARS-Cov-2 infection being mediated by the ACE2 receptor (27). In addition, electron microscopy analysis in representative COVID-19 cases confirmed the presence of virus within AT2 cells (**Figure 1Ci,ii**) and high magnification revealed virions in the proximity of the endoplasmic reticulum (**Figure 1Ciii,iv**) indicating their likely assembly and budding, as well as virions residing in cytoplasmic vesicles (**Figure 1Ciii,v-vi**), implying their transfer and release into the extracellular space.

**Figure 1:**
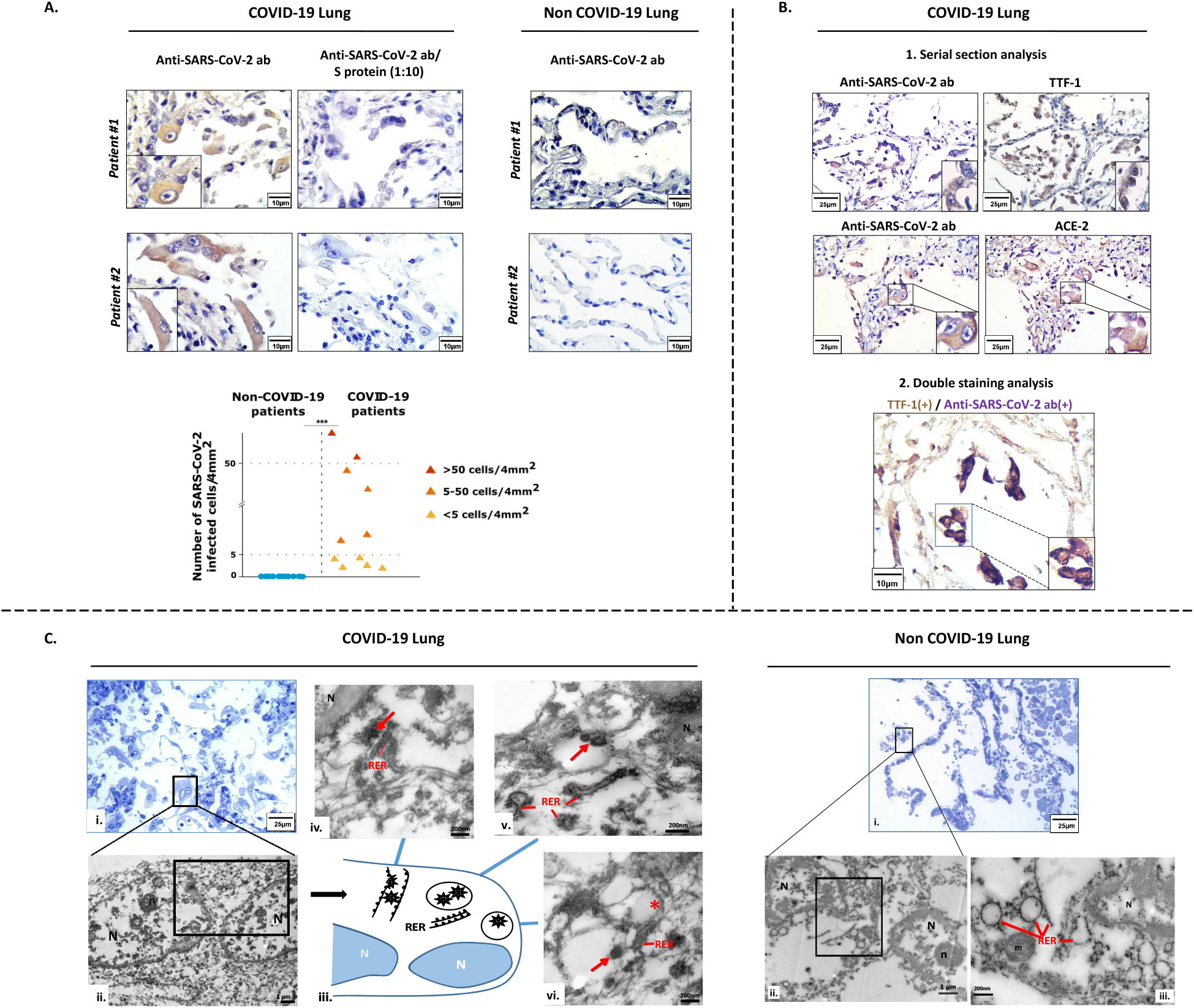
Detection of SARS-CoV-2 in lung cells. **A.** Representative images of SARS-CoV-2 IHC staining in COVID-19 lung tissue. Competition with anti-peptide (S protein) showing specificity of the IHC staining. Representative negative, control IHC staining in non-COVID-19 lung tissues. Graph shows quantification of SARS-CoV-2 staining in the clinical samples (**Suppl Table 1**). **B.** Detection of SARS-CoV-2 in AT2 cells (confirmed by TTF-1 staining) and in ACE-2 expressing cells. Double IHC staining for SARS-CoV-2 and TTF-1. **C.** Detection of SARS-CoV-2 by transmission electron microscopy (TEM) in a representative COVID-19 patient. Presence of SARS-CoV-2 within AT2 cells (**i,ii**) and of virions in the proximity of the endoplasmic reticulum (**iii,iv**) as well as in cytoplasmic vesicles (**iii,v-vi**). Corresponding scale bars are depicted. ICH: immunohistochemistry; AT2: alveolar type 2 cells; ACE2: angiotensin-converting enzyme 2, ***Statistical significant: p<0.001.

### Senescence in SARS-CoV-2 infected cells

A proportion of SARS-CoV-2 infected AT2 cells (range 8 to 21%) displayed a senescent phenotype, with positive staining for SenTraGor and p16^INK4A^ (**Figure 2A-C**) (12,26–28). By contrast lung tissues from age-matched non-COVID-19 cases with analogous co-morbidities (**Suppl Table 1**) showed significantly lower senescence (range 1-2%, p<0.01, Wilcoxon paired non-parametric test) (**Figure 2A-C**), suggesting that SARS-CoV-2 infection may induce senescence.

**Figure 2:**
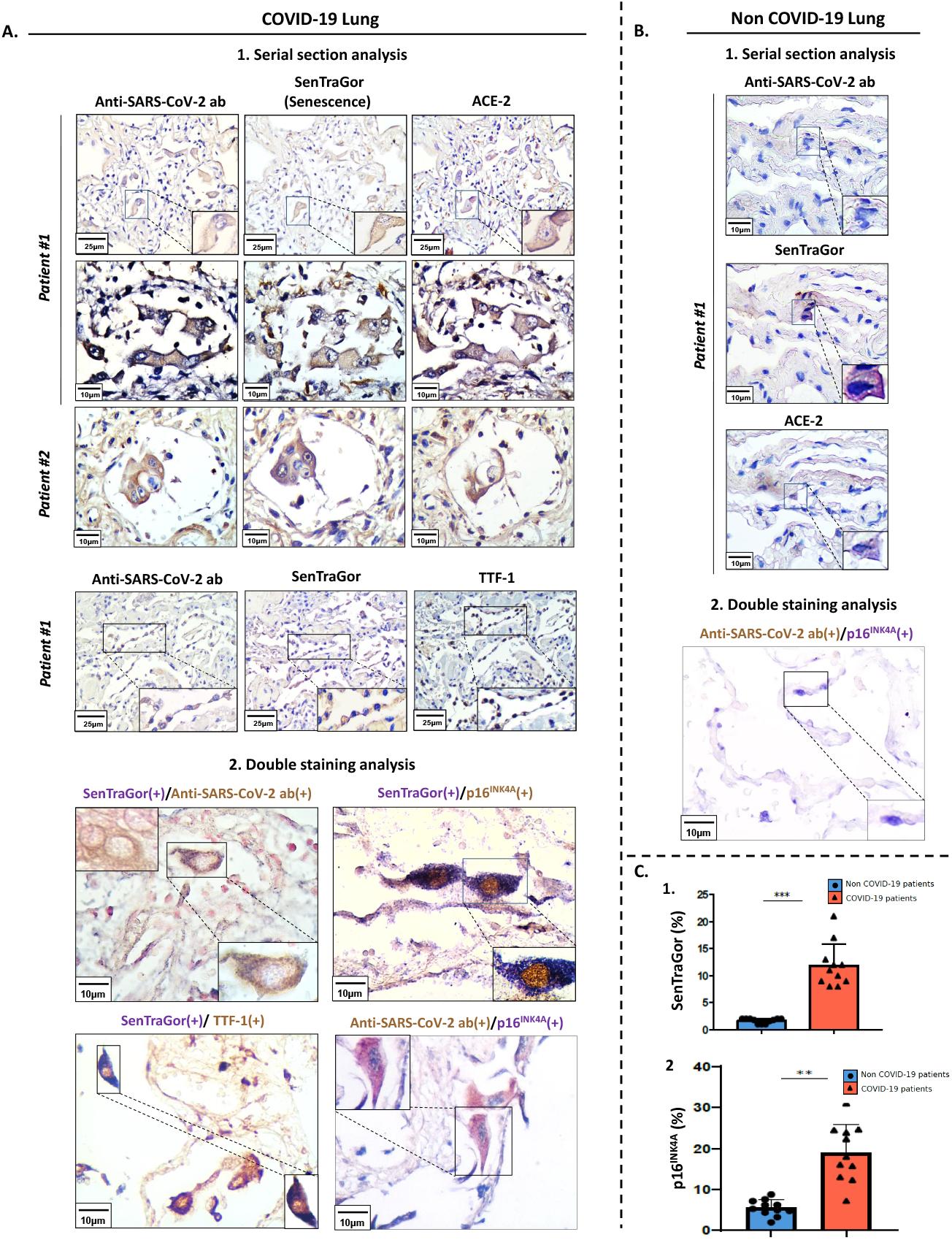
Senescence in SARS-CoV-2 infected cells. **A.** Representative images of SARS-CoV-2, SenTraGor (senescence) and ACE-2 staining in serial sections of COVID-19 lung tissue. Double-immunostaining analysis (**2**) for SARS-CoV-2, SenTraGor (senescence), ACE-2 and p16^INK4A^ in COVID-19 lung tissue. **B.** Representative results from serial staining for SARS-CoV-2, SenTraGor (senescence) and ACE-2, and double-staining experiment for SARS-CoV-2 and p16^INK4A^ in non-COVID-19 lung tissue. **C.** Graphs depicting the increased levels of SenTraGor and p16^INK4A^ in COVID-19 lung tissue. Corresponding scale bars are depicted. Statistical significance: **: p<0.01; ***: p<0.001.

To functionally reproduce our hypothesis we infected Vero cells with a viral strain isolated from a COVID-19 patient. Vero cells is an established cellular system for viral propagation and studies, as apart from their high infectivity to SARS-CoV-2 they are among the few cell lines demonstrating SARS-CoV-2-mediated cytopathic effects, an essential aspect in diagnostics (29,30). Infection was carried out at a low MOI to mimic natural coronavirus infection (31). In line with our hypothesis, the infected cells following an initial surge of cell death reached an equilibrium demonstrating clear evidence of senescence, as compared to the non-infected control cells, 17 days post infection (**Figure 3**). As Vero cells lack p16^INK4A^ (32), the most likely trigger of senescence is DNA damage, as previously reported (12,33). DNA damage measured by γ-H2AX immunostaining, was evident in SARS-CoV-2 infected cells (**Figure 3A4**). It appears that genotoxic stress results from a vicious cycle imposed by the virus in host cells as it high-jacks most intracellular protein machineries (11,34).

**Figure 3.**
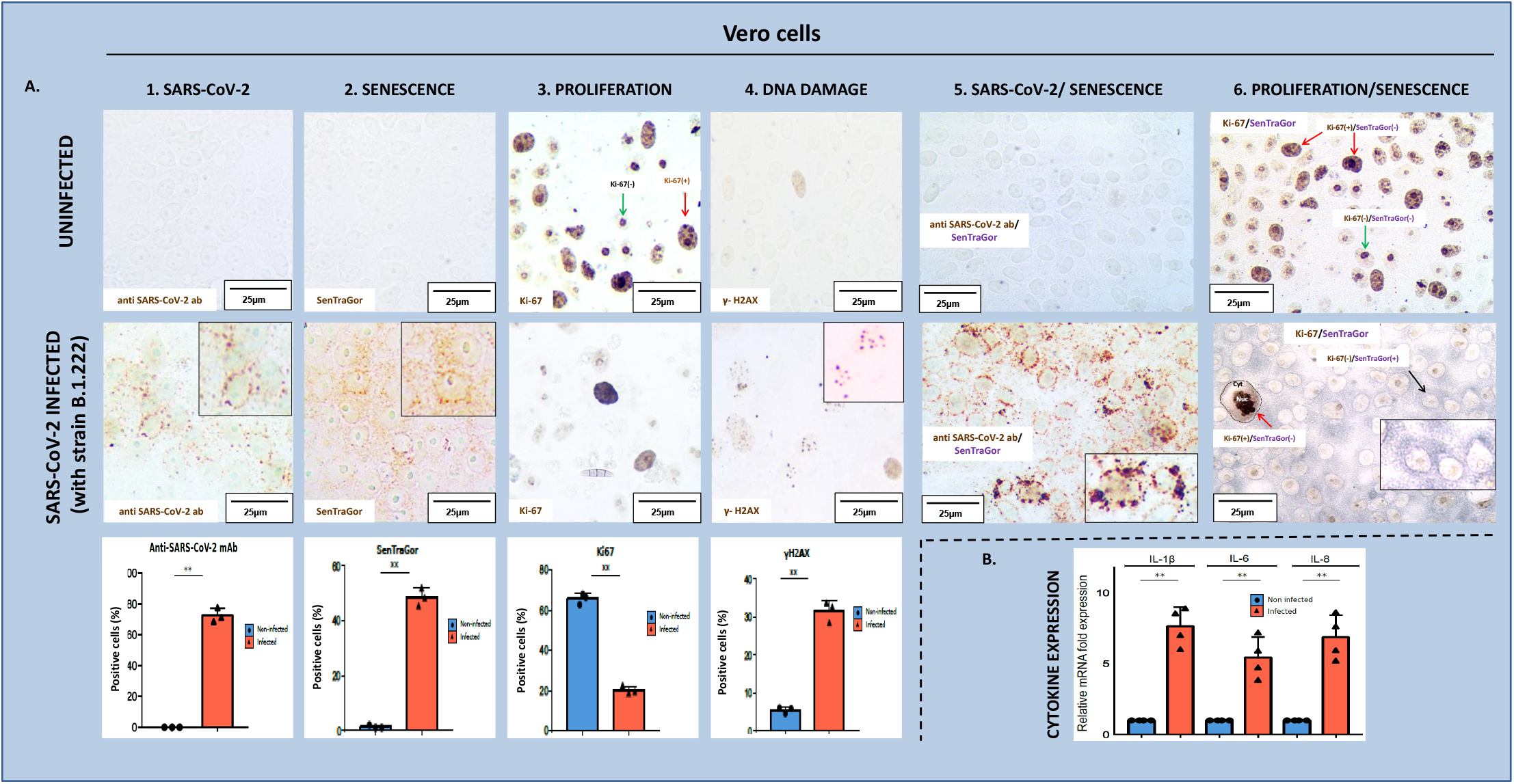
SARS-CoV-2 infection, senescence and SASP expression in Vero cells. **A.** SARS-CoV-2 presence (**1**), senescence induction (**2**), cellular proliferation (**3**) and DNA damage activation (**4**), with corresponding quantitative histograms, in Vero cells with and without SARS-CoV-2 infection. Double IHC staining for SARS-CoV-2 infection/senescence induction (**5**) and senescence induction/cellular proliferation (**6**). **B.** Graph depicting induction of SASP related cytokines following SARS-CoV-2 infection (mRNA expression). **Statistical significant: p<0.01.

### Senescence associated secretory phenotype

We found very high expression of both IL-1β and IL-6 by senescent AT2 cells in the lungs of COVID-19 patients while in the non COVID-19 control cases expression was very low in the few senescent AT2 cells detected (p<0.001) (**Figure 4A-C, Suppl Table 1**). As both cytokines are key components of the “cytokine storm”, our findings suggest putative implication of senescence via SASP in the poor clinical course of COVID-19 patients. Likewise, SARS-CoV-2 senescent Vero cells displayed expression of SASP-related cytokines, as assessed by our recently reported algorithmic assessment of senescence, justifying the *in vivo* findings (**Figure 3B**).

**Figure 4:**
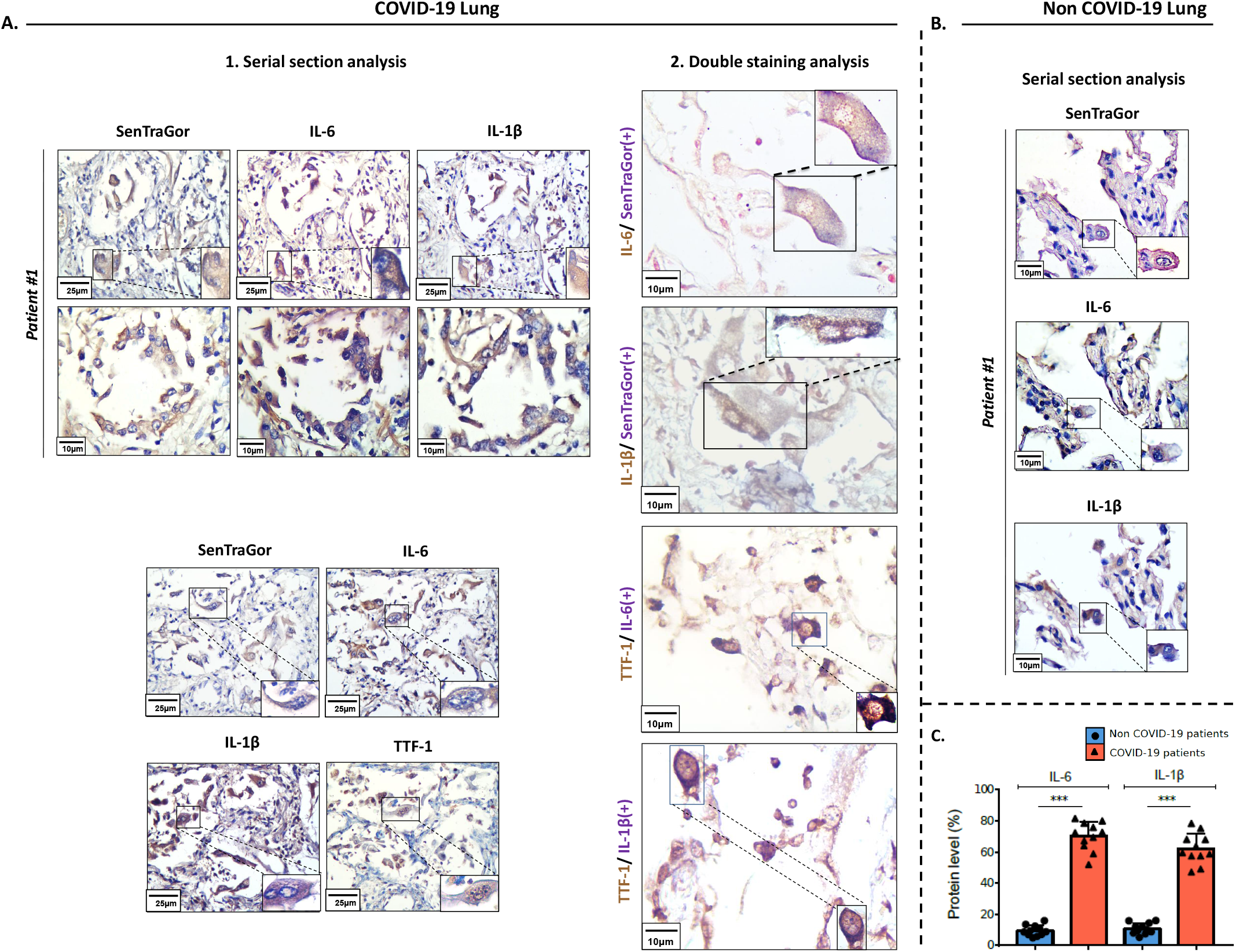
Senescence associated secretory phenotype (SASP) in COVID-19 lung tissues. **A.** Representative staining results (at low and high magnification) of SenTraGor, IL-6, IL-1β and TTF-1 in corresponding serial sections (**1**) and as double immunostaining analysis (**2**) of COVID-19 lung tissue. Original magnification: 400x. **B.** Representative staining results showing absence or minimal levels of SenTraGor, IL-6 and IL-1β in age-matched non-COVID-19 ocntrol samples. Corresponding scale bars are depicted. IL-1β: Interleukin 1β; IL-6: Interleukin-6; ***Statistical significant: p<0.0001.

## Discussion

We have demonstrated the presence of SARS-CoV-2 in AT2 cells of patients who died from COVID-19 using a novel anti-viral antibody and confirmed by electron microscopy. We have shown for the first time that a proportion of SARS-CoV-2-infected AT2 cells acquire senescence features (as demonstrated by significantly increased staining with the novel senescence marker SenTraGor and increased p16^INK4A^). The finding that in age-matched non-COVID-19 cases the percentage of senescent cells was much lower (1-2%) than that of the COVID-19 clinical panel (8-21%), is strongly indicative that SARS-CoV-2 triggers senescence (**Figure 2**). We therefore examined whether cellular infection with SARS-CoV-2 virus (B.1.222 strain) would induce cellular senescence in a susceptible cell line *in vitro* and found that in infected cells there was increased SenTraGor staining, as well as evidence of DNA damage measured by increased γ-H2AX expression. This strongly suggests that SARS-CoV-2 may attach to AT2 cells via ACE2 to infect these cells and through activation of the DNA damage response may induce cellular senescence (34). We also demonstrated that the cells infected with SARS-CoV-2 also show a high degree of expression of IL-1β and IL-6, both components of the SASP and implicated in systemic features of COVID-19 which is associated with a “cytokine storm” (3,4).

Senescent cells are in a state of cell cycle arrest but remain metabolically active and secrete a typical profile of inflammatory proteins known as the senescence-associated secretory phenotype (SASP). SASP components include the proinflammatory cytokines IL-1β and Il-6, which are elevated in COVID-19 patients that have acute respiratory distress syndrome (ARDS) or systemic inflammatory features. The SASP components could induce senescence in nearby cells (paracrine) or may spread senescence systemically (endocrine), thus amplifying this chronic inflammation. It is likely that SARS-CoV-2 spreads from epithelial cells in the lower airways to infect AT2 cells, which express ACE2, and cause local senescence and inflammation in the lung. The virus may then enter the circulation and senescence may subsequently spread systemically to affect other organs, leading to multi-organ failure and death (1,22)

An additional implication relates to the prolonged survival of senescent cells that are infected with the virus, as senescent cells are resistant to apoptosis (12,14). This may allow the virus to be hosted for longer periods compared to other cells with higher cell turnover, exposing its genome to host-mediated editing (35–38). Within this context, we recently reported abundance of the APOBEC enzymes, particularly G and H (RNA editing cytoplasmic variants), which are reported to play a pivotal role in viral RNA editing, in cells undergoing stress-induced senescence (24,39,40). In support to this notion, are the increased APOBEC 3G and 3H expression levels found in the infected Vero cells (**Suppl Figure 3)**. Moreover, by conducting a detailed bioinformatic analysis of 423000 SARS-CoV-2 strains available in the GISAID database, we found that APOBEC signatures seem to potently determine the mutational profile of the SARS-CoV-2 genome (**Suppl Figure 4, Suppl Figure 5).**

A limitation of the study is the small sample size of examined COVID-19 lung autopsies, due to difficulty of getting access to this material. Another limitation due to the nature of the disease is that pathological features, such as senescence, can only be investigated within the context of cadaverous material, which represents the most severe outcome of the spectrum of COVID-19 clinical manifestations. Therefore, evaluation of senescence in less severe conditions is not feasible. Findings have been observed in lung biopsies only and ideally should be also investigated in organs other than the lung.

Overall, SARS-CoV-2 induced senescence justifies the application of senotherapeutics not only as a therapeutic approach for the treatment of COVID-19 patients but also as a putative strategy to restrict mutational events that may favor the emergence of SARS-CoV-2 quasispecies (15,22,41). Senotherapies include senostatics that inhibit components of the cellular senescence pathways and senolytics, which induce senescent cells to become apoptotic (15,22). Several senolytic therapies have been shown to be effective in animal models of accelerated ageing diseases, including COPD, idiopathic pulmonary fibrosis, atherosclerosis and chronic kidney disease (12,15,22). A trial of senolytic therapy in patients with diabetic kidney disease demonstrated a reduction in senescent cells in the skin and reduced circulating SASP proteins, such as IL-1β and IL-6 (42). A clinical trial of a Senolytic compound (F) to inhibit progression to cytokine storm and ARDS in COVID-19 patients has been approved by the US Food and Drug Administration (FDA) and is anticipated to be soon launched (43).

## Acknowledgements

We would like to thank Konstantinos Ntostoglou for his valuable help in preparing the material. We acknowledge support in RNA sequencing by the “The Greek Research Infrastructure for Personalised Medicine (pMED-GR)” (MIS 5002802) which is implemented under the Action “Reinforcement of the Research and Innovation Infrastructure”, funded by the Operational Programme “Competitiveness, Entrepreneurship and Innovation” (NSRF 2014-2020) and co-financed by Greece and the European Union (European Regional Development Fund). This work was supported by the: National Public Investment Program of the Ministry of Development and Investment / General Secretariat for Research and Technology, in the framework of the Flagship Initiative to address SARS-CoV-2 (2020ΣE01300001); Horizon 2020 Marie Sklodowska-Curie training program no. 722729 (SYNTRAIN); Welfare Foundation for Social & Cultural Sciences, Athens, Greece (KIKPE); H. Pappas donation; Hellenic Foundation for Research and Innovation (HFRI) grants no. 775 and 3782 and NKUA-SARG grant 70/3/8916.

## Supplementary Information

### Suppl Figure legends

**Suppl Figure 1: SARS-CoV-2 antibody production and screening selection. A.** Workflow of the procedure for antibody production. **B.** Sequel of screening steps for antibody production and selection. **C.** Final screening step processes leading to the selection of G2 monoclonal antibody (**Suppl Figure 2**).

**Suppl Figure 2:** Graph depicting the structure of G2 antibody as well as the DNA sequences of FRs and CDRs elements of variable regions.

**Suppl Figure 3:** Graph depicting abundance of APOBEC 3G and 3H expression levels in Vero cells by RT-qPCR analysis. *Statistical significant: p<0.05.

**Suppl Figure 4:** APOBEC consensus RNA 2D sequence and structure motifs. **A.** Mutation profile of SARS-CoV-2 genome exhibits APOBEC mutation signatures. Applying bioinformatics analysis, C to U mutations were found to be the most dominant (55%), suggesting an APOBEC driven signature. 56.8% of these mutations were confirmed to exert APOBEC binding characteristics. **B.** Depicts the APOBEC consensus 2D structure image as obtained from Beam software with statistics for the motif as shown in **(C)** and the position of the motif on the 120 nt window as presented in **(D)**. APOBEC average probability per base of being unpaired around a 120 nt region of the most probable C→U site. **E.** The average probability from all C→U sites as obtained from the RNAplfold algorithm is depicted (**i**) while **(ii)** presents the consensus sequence motif relative to the average probability window.

**Suppl Figure 5: Infected senescent cells as a putative source for SARS-CoV-2 quasispecies generation. A. (i)** Schematic layout presenting representative APOBEC sites from the GISAID database analysis that overlap with the C→ U sites of Vero cells after 17 days of infection with the SARS-CoV-2 B.1.222 strain (see also panel **B**). The yellow bars show the frequency of C→U substitutions when observing the GISAID database read counts (green pileups) and with red is the C→U frequency when observing the SARS-CoV-2 genome, 17 days post infection. These representative sites are ranked as highest relatively to the C→U counts as observed from the GISAID database. The genomic co-ordinates of each C→U can be observed at the superimposed SARS-CoV-2 genome. On the left of the graphs is the consensus motif when performing a motif analysis of all the C→U sites of the SARS-CoV-2 genome, 17 days post infection. NGS reads were confirmed in triplicate reads. **(ii)** Graph depicting frequency of nucleotide substitutions that accumulated in the genome of B.1.222 strain following 17 days of infection. **(iii)** Pie chart demonstrating that predominant C→U substitutions (65%) are APOBEC driven. **B.** Additional locations of C→U substitutions observed in the genome of the SARS-CoV-2 progeny after 17 days of infection in Vero cells (relative to panel **A**).

## Material and Methods

### RNA extraction and Reverse-Transcription real-time PCR (RT-qPCR) detection

#### SASP cytokine and APOBEC G and H mRNA analysis

RNA was extracted using the Nucleospin RNA kit (Macherey-Nagel #740955) according to the manufacturer’s instructions. 1 μg RNA was used for cDNA preparation with Primescript™ RT Reagent Kit (Takara #RR037A). RT-qPCR was performed utilizing SYBR Select Master Mix (Life technologies #4472908) on a DNA-Engine-Opticon (MJ-Research) thermal cycler. Primer sequences employed were: *IL-1β* Fw: 5’-GGAAGACAAATTGCATGG-3’, Rv: 5’-CCCAACTGGTACATCAGCAC-3’; *IL-6* Fw: 5’-AGAGGCACTGGCAGAAAAC-3’, Rv: 5’-TGCAGGAACTGGATCAGGAC-3’; *IL-8* Fw: 5’-AGGACAAGAGCCAGGAAGAA-3’, Rv: 5’-ACTGCACCTTCACACAGAGC-3’; *APOBEC3G* Fw: 5’-CCGAGGACCCGAAGGTTAC-3’, Rv: 5’-TCCAACAGTGCTGAAATTCG-3’; *APOBEC3H* Fw: 5’-CGACGGCTTGAAAGGATAGAG-3’, Rv: 5’-TGAGTTGTGTGTTGACGATGA-3’; B2M: β2-microglobulin (reference) gene Fw: 5’-TCTCTGGCTGGATTGGTATCT-3’, Rv: 5’-CAGAATAGGCTGCTGTTCCTATC-3’ (1). Results, averaged from three independent experiments, are presented as n-fold changes after Sars-CoV-2 infection relatively to the non-infected condition, using the 2-ΔΔCT method.

#### Viral RNA detection

RNA was extracted using the NucleoSpin Virus RNA purification kit (Macherey-Nagel #740.983) according to the manufacturer’s instructions. RT-qPCR was performed utilizing the One Step PrimeScript III RT-PCR Kit (Takara # RR601B) on a Rotor-Gene Q 6000 (Qiagen) thermal cycler following the manufacturer’s instructions and using the CDC N-gene directed primers [**https://www.cdc.gov/coronavirus/2019-ncov/lab/rt-pcr-panel-primer-probes.html**],

### Anti-SARS-COV-2 antibodies

#### Generation

A series of monoclonal antibodies against SARS-CoV2 spike protein were produced according to a modified method of Koehler and Milstein (Koehler and Milstein, 1975). Briefly, twelve BALB/c mice of 5 weeks of age were immunized intraperitoneally (i.p.) with 25μg of SARS-Cov2 protein (Trenzyme GmbH, Germany). All immunization and animal handling were in accordance with animal care guidelines as specified in EU Directive 2010/63/EU. After 5 cycles of immunization, mice were sacrificed, spleenocytes were collected and fused with P3X63Ag8.653 (ATCC® CRL1580™) following a modified method of Koehler and Milstein. Positive clones and antibody specificity were determined through extensive immunosorbent assays. Four clones, namely 479-S1, 480-S2, 481-S3 and 482-S4 are under patent application (**Gorgoulis V.G., Vassilakos D. and Kastrinakis N. (*2020*) GR patent application no: 22-0003846810**).

#### RNA sequence determination and amino acid prediction

RNA was collected from biological duplicates of generated hybridomas as described elsewhere (2). RNA samples were processed according to manufacturer’s instructions, using the following kits: NEBNext® Poly(A) mRNA Magnetic Isolation Module (E7490S), NEBNext^®^ Multiplex Oligos for Illumina^®^ (Index Primers Set 1, NEB7335) and NEBNext^®^ Ultra™ II Directional RNA Library Prep with Sample Purification Beads (E7765S). After successful QC (RNA 6000 Nano bioanalyzer, Agilent) and quantity measurements (Qubit™ RNA HS Assay Kit, Thermofisher), 1ug was used for mRNA selection, cDNA construction, adaptor ligation and PCR amplification (11 cycles), according to the manufacturer’s protocol: (https://international.neb.com/products/e7760-nebnext-ultra-ii-directional-rna-library-prep-kit-for-illumina#Product%20Information). The 479-G2-ATCACG index from NEB E7335 was used. The final libraries were analyzed with Agilent High Sensitivity DNA Kit on an Agilent bioanalyzer, quantitated (Qubit dsDNA HS Assay Kit, Thermofisher) and, after multiplexing, were run using a NextSeq 500/550 Mid Output Kit v2.5 (150 cycles), paired end mode on a NextSeq550 (Illumina) at final concentration 1,3pM with 1% PhiX Control v3.

Fastq files were demultiplexed with Flexbar (3). Quality control of the Fastq files was assessed with FastQC tools (4). Adapter sequences were removed with Cutadapt program (5) with the following parameters: quality trimming was set to 20 and the minimum allowed nucleotide length after trimming was 20 nucleotides using --pair-filter=any to apply the filters to both paired reads. A two way alignment mode was followed to identify the antibody clone. More precisely alignments were performed with Bowtie2 (6) with parameters set as following: -D 20 -R 3 -N 1 -L 20 -i S,1,0.50 –no-mixed --no-discordant against an index made from IMGT database http://www.imgt.org/ having downloaded all mouse and human IG genes. Also this mode of alignments was executed for quality control and visualization of the aligned reads spanning the IG gene segments on the genome browser. The second mode refers to the determination and reconstruction of the clones. This was performed with MiXCR suite (7). At first, alignments against the IG repertoire were performed with kaligner and visualization of alignments was assessed. It was observed that the use of kaligner gave better results with higher clone hits regarding the VH and VL segments. Full assembly of the clones was performed. A full report of the number of reads and assembly of CDR and FR clones is provided in clones479_S1kalign.txt. The clones with the highest number of reads and coverage across the V,D,J segments were considered. The reported matched sequences were also checked with IgBlast tool https://www.ncbi.nlm.nih.gov/igblast/. In addition, after the assembly of the amino acid reconstruction of the FR and CDR regions of the full variable fragment for both the Heavy and Light antibody chains, a 3D visualization was also determined via folding the V protein fragment with iTassser suite (8). The above analysis has been extensively described in **Gorgoulis VG, Vassilakos D and Kastrinakis N. (*2020*) GR patent application no: 22-0003846810**.

### Immunocytochemistry (ICC)-Immunohistochemistry (IHC)

#### Method

ICC and IHC were performed according to previous published protocols (9). In brief, 3 μm thick sections from formalin-fixed paraffin embedded (FFPE) lung tissues were employed. Antigen retrieval was heat-mediated in 10 mM citric acid (pH 6.0) for 15 minutes. The following primary antibodies were applied: i) the anti-SARS-CoV-2 (G2) monoclonal antibody (dilution 1:300), ii) anti-ACE-2 [Rabbit polyclonal antibody Abcam, Cat.no: ab15348 (dilution 1:200)], iii) anti-TTF-1 [rat monoclonal antibody Dako, Clone 8G7G3/1, Cat.no: M3575 (Ready-to-Use)], iv) anti-CD68 [mouse monoclonal antibody Dako, Clone PG-M1, Cat.no: M0876 (dilution 1:50)] and v) anti-p16^INK4A^ [mouse monoclonal antibody Santa Cruz, clone: F-12, Cat.no.:sc-1661. (dilution 1:100)], vi) IL-1β [Rabbit polyclonal antibody Abcam, Cat.no: ab2105 (dilution 1:150)] and vii) IL-6 [mouse monoclonal antibody R&D systems, clone: Clone: 6708, Cat.no:MAB206 (dilution 1:100)], all overnight at 4°C. Development of the signal was achieved using the Novolink Polymer Detection System (Cat.no: RE7150-K, Leica Biosystems). Specimens were counterstained with hematoxylin.

#### Negative Controls for the anti-SARS-CoV-2 (G2) monoclonal antibody

*i) Biological*, comprising previously published and new lung tissue samples from a cohort of 50 cases that underwent surgery prior to COVID-19 outbreak. *ii) Technical*: a. Omission of the G2 primary monoclonal antibody, b. Blocking of the G2 primary monoclonal antibody using the corresponding S-protein (Cat.no.P2020-029, Trenzyme) in a 1:10 (G2/Spike protein) ratio and c. Two slides per case were employed for each staining or control experiment.

#### Evaluation of G2 staining

Cells were considered positive irrespective of the staining intensity. Two different semi-quantitative IHC evaluation approaches, previously described were adopted (10,11) According to the first, the number of G2 positive cells per 4mm^2^ was encountered and scored according to the following criteria: (+) for positive staining in<5 cells per 4 mm^2^, (+) for positive staining in 5–50 cells per 4mm^2^ and (+++) for positive staining in >50 cells per 4 mm^2^ (10). Regarding the second one, the number of G2 positive cells per whole slide was estimated and subsequent scores were assessed: (+) between one and five positive cells per whole slide (scattered cells), (++) more than five cells per whole slide but no foci (isolated cells) and (+++) more than 10 cells in one × 20 field (with foci) (11). For IL-6 and IL-1b, the percentage of immunopositive cells was encountered (12). Evaluations were performed blindly by four experienced pathologists (KE, PF, CK and VG) and intra-observer variability was minimal (p≤0.05).

### Bioinformatic analysis for identification of mutational signatures in the SARS-CoV-2 genome

#### Screening for mutational signatures in the SARS-CoV-2 genome

To investigate the mutational properties on the SARS-CoV-2 genome we downloaded from GISAID database (**https://www.gisaid.org/**) 423.000 available strains that were distributed globally. These strains were aligned with the Wuhan first assembly NC_045512, obtained from NCBI (**https://www.ncbi.nlm.nih.gov/sars-cov-2/**), with Bowtie aligner (13) using the following command:

/bowtie2-2.4.2-sra-linux-x86_64/bowtie2-align-s --wrapper basic-0 -x Covncbiref -p 4 -D 20 -R 3 -N 1 -L 20 -i S,1,0.50 -f allCov19.fa

In order to identify the mutations we have created an “in-house” script using *calmd* function from SAMtools (14), based on the analysis of deciphering mutations from the proteome occupancy profile study (15). We applied the following commands for minus and reverse stranded reads:

*samtools sort accepted_hits.bam -o accepted_hitsort.bam*
*samtools rmdup -s accepted_hitsort.bam rmdupsorted.bam*
*#forward library*
*samtools view -h -f 0×0010 rmdupsorted.bam | samtools calmd -S -*
*~/Desktop/Bioinformatics/NCBI.fa /dev/stdin |*
*/media/covid_meth/deademination_covid19/get_edit_stat.pl ‘-’ >*
*Mapping_editStatus.bed*
*samtools view -h -F 0×0010 rmdupsorted.bam | samtools calmd -S -*
*~/Desktop/Bioinformatics/NCBI.fa /dev/stdin |*
*/media/covid_meth/deademination_covid19/get_edit_stat.pl* ‘+’ >>
*Mapping_editStatus.bed*
*#reverse library*
*samtools view -h -F 0×0010 rmdupsorted.bam | samtools calmd -S -*
*~/Desktop/Bioinformatics/GRCh37/hg19.fa /dev/stdin |*
*/media//covid_meth/deademination_covid19/get_edit_stat.pl ‘-’ >*
*Mapping_editStatus2.bed*
*samtools view -h -f 0×0010 rmdupsorted.bam | samtools calmd -S -*
*~/Desktop/Bioinformatics/GRCh37/hg19.fa /dev/stdin |*
*/media//covid_meth/deademination_covid19/get_edit_stat.pl ‘+’ >>*
*Mapping_editStatus2.bed*

The scripts bellow were used in order to determine the counts per type of mutation and filter for C→U or G→A mutations in respect with the strand orientation of the alignments. Bedtools (16) have also been used to obtain the fasta sequences and the windows around the C→U sites.

#sort reads
sort -k1,1 -k2,2n -k3,3n*Mapping_editStatus2.bed*| uniq -c >
Mapp_editstat_APOBECcounts.bed sort -k1,1 -k2,2n -k3,3n*Mapping_editStatus2.bed*|
uniq -c > Mapp_editstat_APOBECcounts.bed sed -i ‘s/^ *//g’
Mapp_editstat_APOBECcounts.bed
#obtain the fasta
fastaFromBed -s -fi ~/Desktop/Bioinformatics/GRCh37/hg19.fa -
bedMapp_editstat_APOBECcounts.bed -tab -foAPOBEC_counts1fa.bed
#get mutation type C→ U or A→G
perl fixmutstat.pl APOBEC_counts1fa.bed APOBEC_counts1facorrect

Based on the filtered candidate sites with a frequency of mutations above than 5 reads we have obtained windows of ±60 nucleotides and folded the RNA sequences from these regions with Vienna RNA fold algorithm (17) to determine the RNA 2D structure. SHAPE reactivities from SHAPE-seq data (18) were used to guide the RNA folding.

*RNAfold --noPS --shape=forViennatest.SHAPE.txt –shapeConversion=S -g < forViennatest..fa*
*>> test.txt*.

To decipher the candidate motifs we counted the frequency of letters ±5 nucluteotides from the most frequent deademinated nucleotide. The frequency for each letter was determined via a perl script which extracts all possible *k-mers* and their frequencies. Next these *k-mers*, based on their frequency, were plotted with Web-logo motifs (19). In addition position-weight matrices (PWM) for each letter around the deadiminated RNA nucleotide were extracted.

Our analysis on motifs and RNA structure for choosing the candidate APOBEC sites based on publicly available known studies (20–22) that also demonstrate similar characteristics regarding the motif specific APOBEC signature and RNA structure. From our analysis we determined a CCT/A enrichment around regions of open hairpin structures agreeing with the results from the literature.

#### Verification of APOBEC specific motifs by applying machine learning

To filter and obtain scores for each APOBEC specific candidate site we have also applied a machine learning scheme using convolutional neural networks having as input the sequence and RNA structure around the candidate strongest APOBEC sites with high frequency that also demonstrate a high potential for APOBEC binding.

#### APOBEC consensus RNA 2D sequence and structure motifs

To determine the consensus RNA structure properties we have used the Vienna RNA folding output dot bracket notation, which performed the folding based on the icSHAPE reactivities as input to BEAM program (23). In addition, the binding sites from (19) have been used to decipher the structure for the hg19. The structure properties for the APOBEC sites on the hg19 have been folded using SHAPE (24) and DMS (25) data to guide the RNA folding. Secondary RNA structure motifs regarding the hg19 have been determined using BEAM software. Beam software was used as follows:

First the dot bracket notation is translated in a 24 letter language for structure called Bear:

*java -jar encoder.jar APOBEC.db APOBEC.fb*
*java -jar /$basepath3/BEAM_release_1.5.1.jar -f APOBEC.fb -w 15 -W 30 -M 5*

Next, then consensus motifs are extracted, using as maximum threshold to output the top 5 motifs with maximum width of motif set to 30 nucleotides.

### APOBEC average probability per base of being unpaired around a 120 nt region of the most probable C→U site

Furthermore RNAplfold has been used to extract the probabilities per base of being unpaired having the following parameters

*RNAplfold -W 30 -L 15 -u 1 --shape=forViennatest.SHAPE.txt --shapeMethod=D --shapeConversion=O -g <forViennatest..fa >> test.txt*

The output per sequence is a _*lunp* file where from these files the smoothed geometric mean per base pair is extracted and plotted having the candidate deaminated site, located in the center of the 120 nucleotide window. The plots are done in R using fit and polygon functions from the standard R bioconductor packages. The confidence intervals per base are calculated as 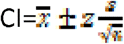 where z =0.95 % confidence, s=standard deviation and *n* are the number of sequences ~3500.

In order to accomplish docking of the APOBEC with the RNA substrates we have used SimRNA suite (26) to determine the RNA 3D structure properties of the SARS-CoV-2 RNA around the candidate C→U sites. DARS-RNP potential (27) has been used using PDB files from crystallographic data for APOBEC3G with PDB code 6bux and 6k3j (28) from the PDB database (**https://www.rcsb.org/)** for docking the RNAs as obtained from SimRNA with APOBEC. Scores were ranked and the RNAs with docking scores higher than 1 standard deviation over the mean were used for the extraction of consensus motifs both in terms of structure and sequence. Regarding the sequence motif a window of ±10 nt around the high docking score was obtained and according to the *k-mer* distribution PWM matrices are extracted and plotted with web-logo.

The SimRNA commands to extract the 3D RNA structure are the following:

./SimRNA -s 3D_testVf.fa -c config2.dat -S 3D_test.struct -o tRNAs python2 trafl_extract_lowestE_frame.py tRNAs.trafl”
./SimRNA_trafl2pdbs tRNAs-000001.pdb tRNAs_minE.trafl : AA” perl configpdb2.pl tRNAs_minE-000001_AA.pdb tRNAs.config”

The configuration file for the 3D simulations is set as:

*NUMBER_OF_ITERATIONS 160000*
*TRA_ WRITE_IN_EVERY_N_ITERATIONS 16000*
*INIT_TEMP 1.35*
*FINAL_TEMP 0.90*
*BONDS_WEIGHT 1.0*
*ANGLES_WEIGHT 1.0*
*TORS_ANGLES_ WEIGHT 0.0*
*ETA_ THETA_ WEIGHT 0.40*
*SECOND_STRC_RESTRAINTS_ WEIGHT 1.0*
*FRACTION_OF_NITROGEN_ATOM_MOVES 0.10*
*FRACTION_OF_ONE_ATOM_MOVES 0.45*
*FRACTION_OF_TWO_ATOMS_MOVES 0.44*
*FRACTION_OF_FRAGMENT_MOVES 0.01*

In order to visualize the properties that might determine the binding of APOBEC, the power of integrated gradients tools (29) were used to obtain the motifs. Our analysis is based upon an already developed method DeepRipe (30) adding an extra module to also incorporate the RNA structure information. The classifier has been trained to distinguish such motifs which are characteristic for APOBEC binding.

