## Supplemental Figure 1 for "SARS-CoV-2 infects lung epithelial cells and induces senescence and an inflammatory response in patients with severe COVID-19"

### Slide 1
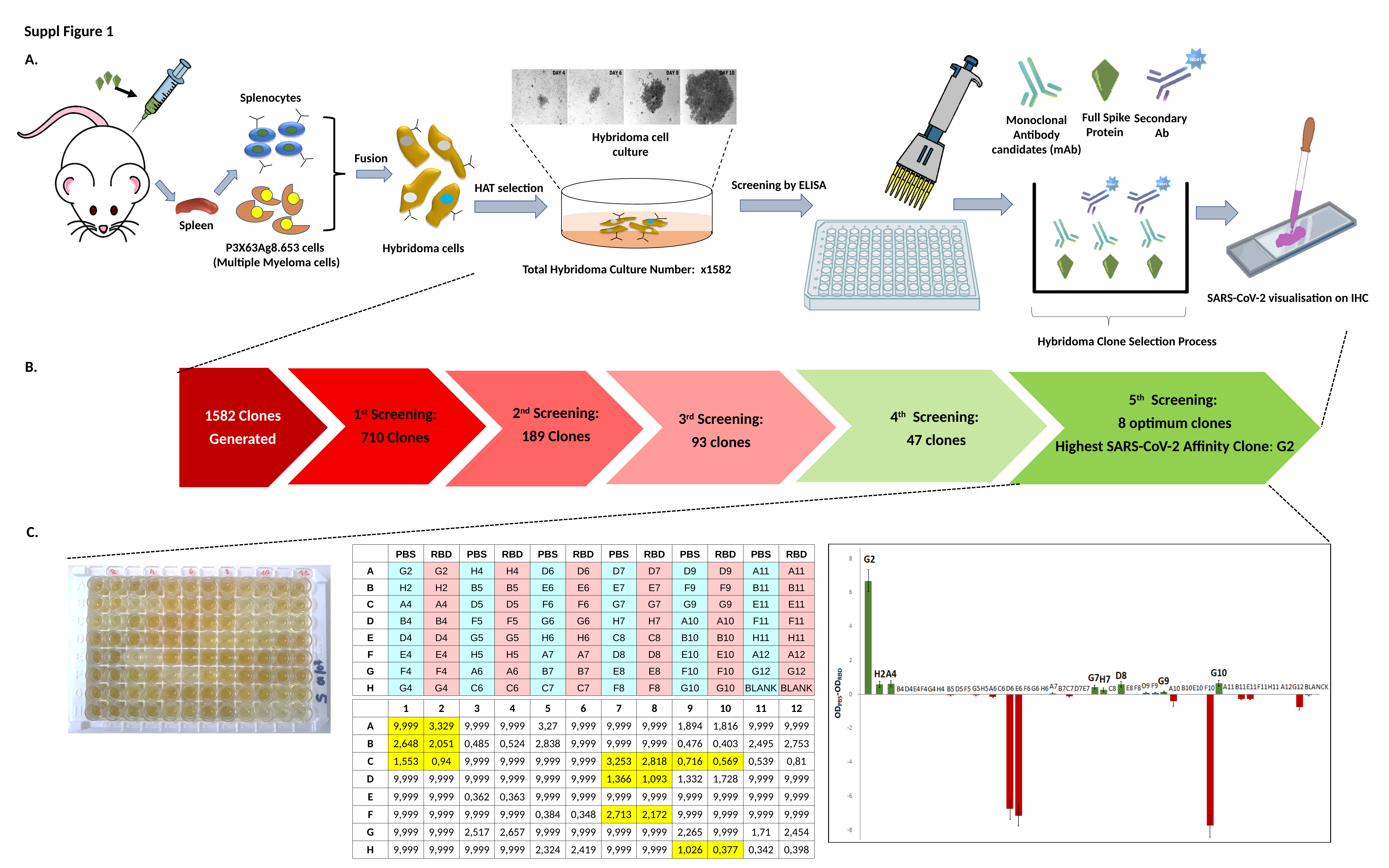

Suppl Figure 1
A.
Splenocytes
Fusion
P3X63Ag8.653 cells
(Multiple Myeloma cells)
Full Spike Protein
Secondary
Ab
Monoclonal Antibody candidates (mAb)
Hybridoma cell culture
Screening by ELISA
HAT selection
Spleen
Hybridoma cells
Total Hybridoma Culture Number: x1582
SARS-CoV-2 visualisation on IHC
Hybridoma Clone Selection Process
B.
2^nd^ Screening:
189 Clones
1582 Clones
Generated
1^st^ Screening:
710 Clones
4^th^ Screening:
47 clones
3^rd^ Screening:
93 clones
5^th^ Screening:
8 optimum clones
Highest SARS-CoV-2 Affinity Clone: G2
C.
| | PBS | RBD | PBS | RBD | PBS | RBD | PBS | RBD | PBS | RBD | PBS | RBD |
| --- | --- | --- | --- | --- | --- | --- | --- | --- | --- | --- | --- | --- |
| A | G2 | G2 | H4 | H4 | D6 | D6 | D7 | D7 | D9 | D9 | A11 | A11 |
| B | H2 | H2 | B5 | B5 | E6 | E6 | E7 | E7 | F9 | F9 | B11 | B11 |
| C | A4 | A4 | D5 | D5 | F6 | F6 | G7 | G7 | G9 | G9 | E11 | E11 |
| D | B4 | B4 | F5 | F5 | G6 | G6 | H7 | H7 | A10 | A10 | F11 | F11 |
| E | D4 | D4 | G5 | G5 | H6 | H6 | C8 | C8 | B10 | B10 | H11 | H11 |
| F | E4 | E4 | H5 | H5 | A7 | A7 | D8 | D8 | E10 | E10 | A12 | A12 |
| G | F4 | F4 | A6 | A6 | B7 | B7 | E8 | E8 | F10 | F10 | G12 | G12 |
| H | G4 | G4 | C6 | C6 | C7 | C7 | F8 | F8 | G10 | G10 | BLANK | BLANK |
| | 1 | 2 | 3 | 4 | 5 | 6 | 7 | 8 | 9 | 10 | 11 | 12 |
| --- | --- | --- | --- | --- | --- | --- | --- | --- | --- | --- | --- | --- |
| A | 9,999 | 3,329 | 9,999 | 9,999 | 3,27 | 9,999 | 9,999 | 9,999 | 1,894 | 1,816 | 9,999 | 9,999 |
| B | 2,648 | 2,051 | 0,485 | 0,524 | 2,838 | 9,999 | 9,999 | 9,999 | 0,476 | 0,403 | 2,495 | 2,753 |
| C | 1,553 | 0,94 | 9,999 | 9,999 | 9,999 | 9,999 | 3,253 | 2,818 | 0,716 | 0,569 | 0,539 | 0,81 |
| D | 9,999 | 9,999 | 9,999 | 9,999 | 9,999 | 9,999 | 1,366 | 1,093 | 1,332 | 1,728 | 9,999 | 9,999 |
| E | 9,999 | 9,999 | 0,362 | 0,363 | 9,999 | 9,999 | 9,999 | 9,999 | 9,999 | 9,999 | 9,999 | 9,999 |
| F | 9,999 | 9,999 | 9,999 | 9,999 | 0,384 | 0,348 | 2,713 | 2,172 | 9,999 | 9,999 | 9,999 | 9,999 |
| G | 9,999 | 9,999 | 2,517 | 2,657 | 9,999 | 9,999 | 9,999 | 9,999 | 2,265 | 9,999 | 1,71 | 2,454 |
| H | 9,999 | 9,999 | 9,999 | 9,999 | 2,324 | 2,419 | 9,999 | 9,999 | 1,026 | 0,377 | 0,342 | 0,398 |
