## Supplemental Figure 2 for "SARS-CoV-2 infects lung epithelial cells and induces senescence and an inflammatory response in patients with severe COVID-19"

### Slide 1
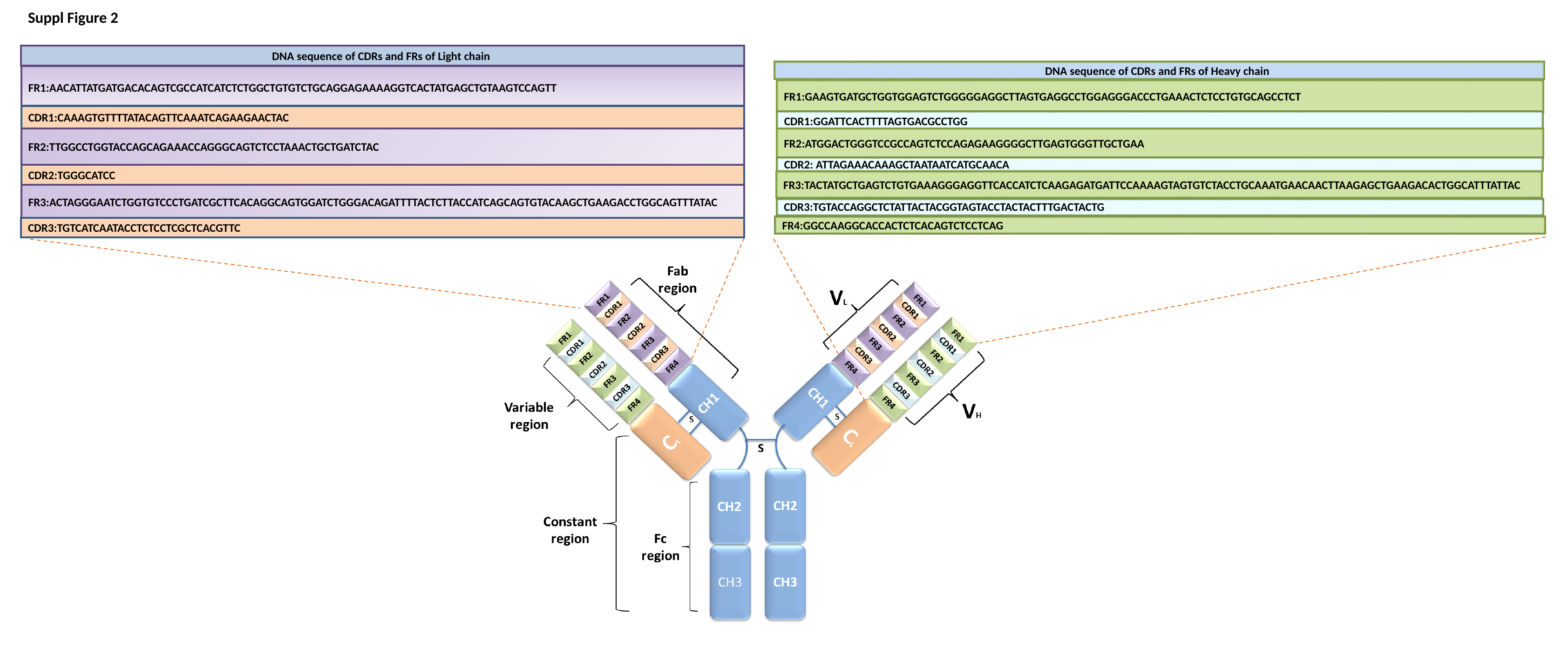

Suppl Figure 2
DNA sequence of CDRs and FRs of Light chain
FR1:AACATTATGATGACACAGTCGCCATCATCTCTGGCTGTGTCTGCAGGAGAAAAGGTCACTATGAGCTGTAAGTCCAGTT
CDR1:CAAAGTGTTTTATACAGTTCAAATCAGAAGAACTAC
FR2:TTGGCCTGGTACCAGCAGAAACCAGGGCAGTCTCCTAAACTGCTGATCTAC
CDR2:TGGGCATCC
FR3:ACTAGGGAATCTGGTGTCCCTGATCGCTTCACAGGCAGTGGATCTGGGACAGATTTTACTCTTACCATCAGCAGTGTACAAGCTGAAGACCTGGCAGTTTATAC
CDR3:TGTCATCAATACCTCTCCTCGCTCACGTTC
DNA sequence of CDRs and FRs of Heavy chain
FR1:GAAGTGATGCTGGTGGAGTCTGGGGGAGGCTTAGTGAGGCCTGGAGGGACCCTGAAACTCTCCTGTGCAGCCTCT
CDR1:GGATTCACTTTTAGTGACGCCTGG
FR2:ATGGACTGGGTCCGCCAGTCTCCAGAGAAGGGGCTTGAGTGGGTTGCTGAA
CDR2: ATTAGAAACAAAGCTAATAATCATGCAACA
FR3:TACTATGCTGAGTCTGTGAAAGGGAGGTTCACCATCTCAAGAGATGATTCCAAAAGTAGTGTCTACCTGCAAATGAACAACTTAAGAGCTGAAGACACTGGCATTTATTAC
CDR3:TGTACCAGGCTCTATTACTACGGTAGTACCTACTACTTTGACTACTG
FR4:GGCCAAGGCACCACTCTCACAGTCTCCTCAG
