## Supplemental Figure 3 for "SARS-CoV-2 infects lung epithelial cells and induces senescence and an inflammatory response in patients with severe COVID-19"

### Slide 1
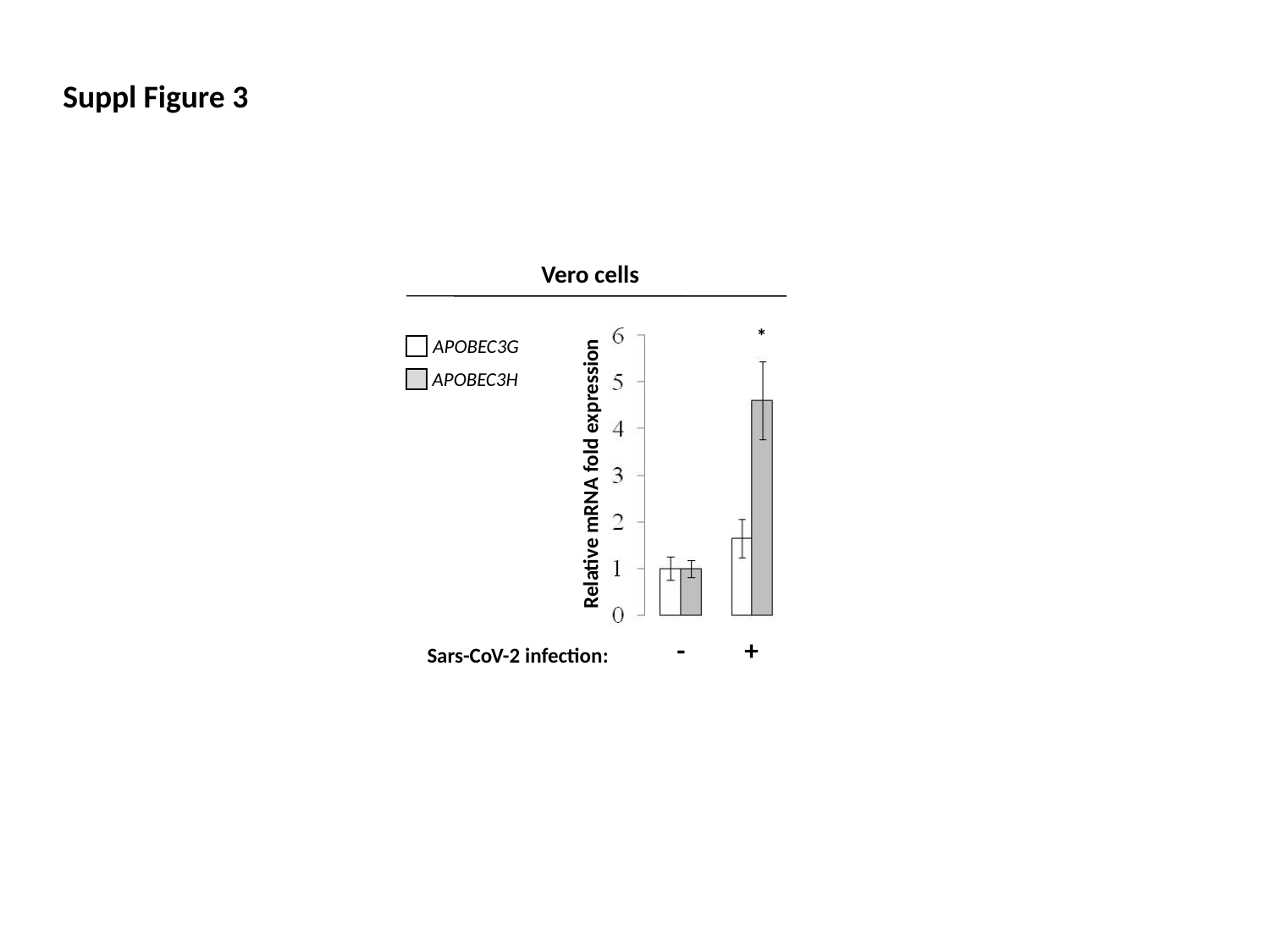

Suppl Figure 3
Vero cells
*
APOBEC3G
APOBEC3H
Relative mRNA fold expression
-
+
Sars-CoV-2 infection:
