## Supplementary figures and images for "SARS-CoV-2 infects lung epithelial cells and induces senescence and an inflammatory response in patients with severe COVID-19"

### Supplemental Figure 4

#### Slide 1
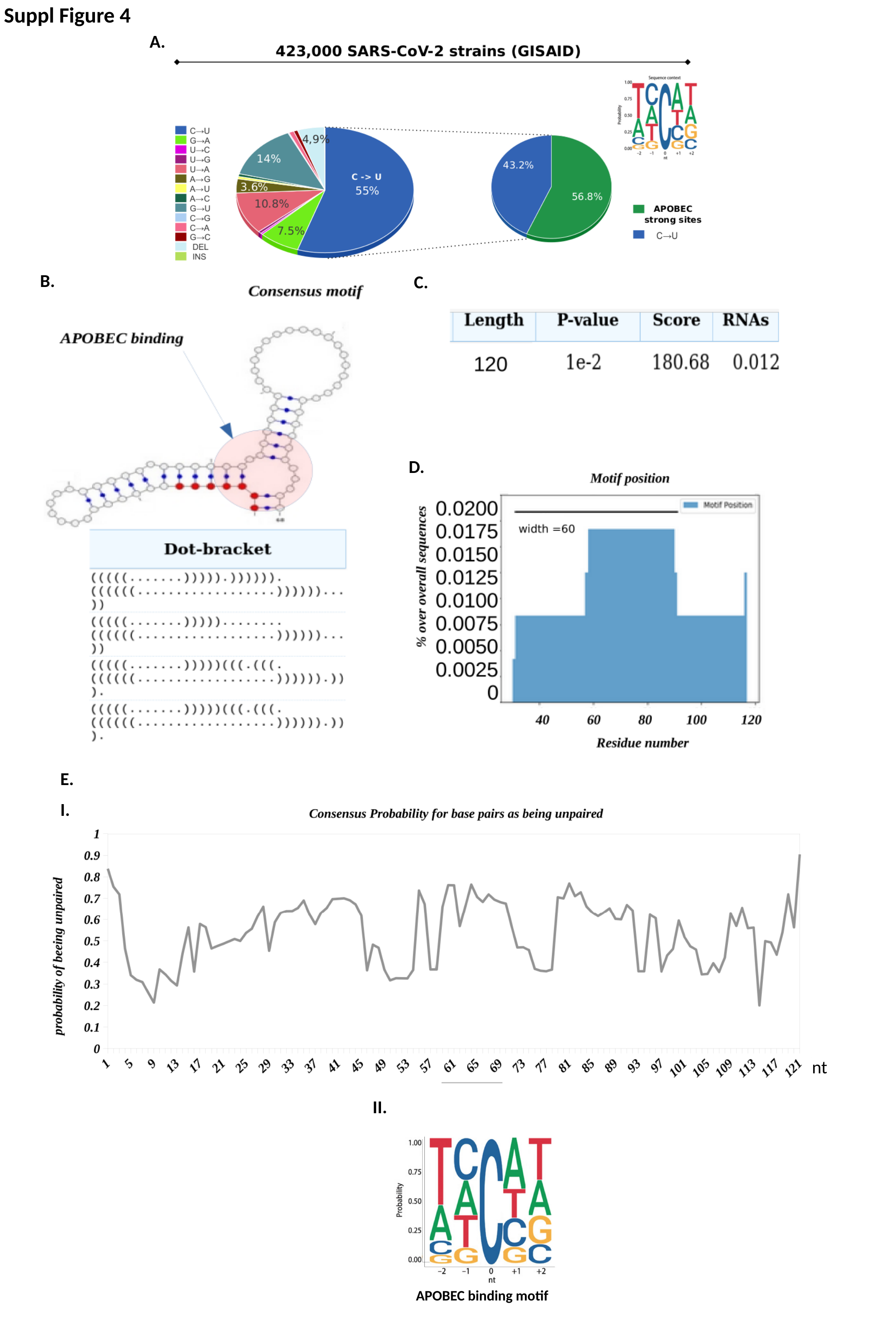

Suppl Figure 4
A.
B.
C.
D.
E.
I.
nt
II.
APOBEC binding motif
