## Supplemental Figure 5 for "SARS-CoV-2 infects lung epithelial cells and induces senescence and an inflammatory response in patients with severe COVID-19"

### Slide 1
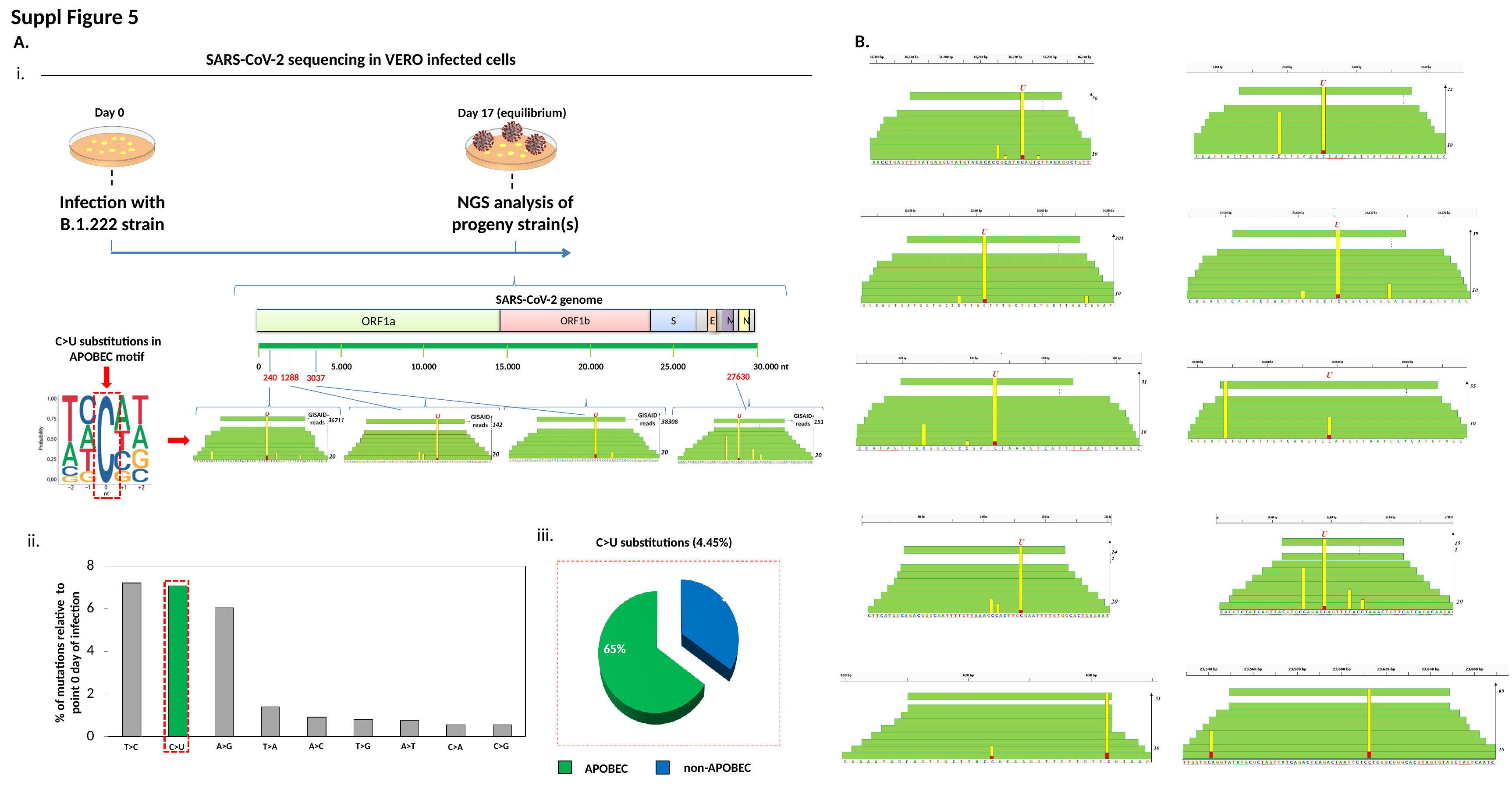

Suppl Figure 5
B.
A.
SARS-CoV-2 sequencing in VERO infected cells
i.
Day 0
Day 17 (equilibrium)
Infection with B.1.222 strain
NGS analysis of progeny strain(s)
SARS-CoV-2 genome
ORF1a
ORF1b
S
M
N
E
C>U substitutions in
APOBEC motif
10.000
15.000
0
5.000
20.000
25.000
30.000 nt
27630
240
1288
3037
iii.
ii.
C>U substitutions (4.45%)
% of mutations relative to point 0 day of infection
A>G
A>C
T>G
A>T
C>G
T>A
T>C
C>U
C>A
35%
65%
non-APOBEC
APOBEC
